# S2Tag, a novel affinity tag for the capture and immobilization of coiled-coil proteins: application to the study of human β-cardiac myosin

**DOI:** 10.1101/2025.05.01.651501

**Authors:** Bipasha Barua, Robert C. Cail, Yale E. Goldman, E. Michael Ostap, Donald A. Winkelmann

**Author notes:** Corresponding author: Donald A. Winkelmann, Department of Pathology and Laboratory Medicine, Robert Wood Johnson Medical School, Rutgers University, Piscataway, NJ 08854.

## Abstract

Single molecule and ensemble motility assays are powerful tools for investigating myosin activity. A key requirement for the quality and reproducibility of the data obtained with these methods is the mode of attachment of myosin to assay surfaces. We previously characterized the ability of a monoclonal antibody (10F12.3) to tether skeletal muscle myosin to nitrocellulose coated glass. Here, we identify the 11 amino-acid epitope (S2Tag) recognized by 10F12.3 in the coiled-coil S2 domain of myosin. To test the transferability of S2Tag, we inserted it into a wild-type β-cardiac myosin construct (WT-βCM) and evaluated its mechanochemistry. WT-βCM immobilized via S2Tag robustly propelled actin filaments in gliding assays and showed single-molecule actin displacements and attachment kinetics by optical trapping. Thus, the antibody attachment is effective for ensemble and single molecule assays. We inserted the S2Tag into a βCM construct containing a penetrant mutation (S532P-βCM) that causes dilated cardiomyopathy. Inclusion of S2Tag enabled quantitative mixed-motor gliding filament assays with WT-βCM. The analysis shows the S532P mutation results in a 60% decrease in gliding speed, yet the motor seems to produce the same force as WT-βCM. Importantly, S2Tag is a useful new tool for affinity capture of alpha-helical coiled coil proteins.

## Introduction

Myosins are a diverse group of motor proteins that play crucial roles in many cellular processes, including cell migration, cell division, intracellular transport, and muscle contraction (1). Assays that measure motor activity have revealed substantial kinetic and motile diversity among the myosin paralogs, leading to a better understanding of their mechanisms, cell functions, and how mutations cause disease. Notably, *in vitro* actin gliding, single-molecule optical trapping, and fluorescence tracking assays have been essential for revealing mechanochemical mechanisms (2-4).

Many *in vitro* assays require attachment of myosin to the surface of glass coverslips, beads, or quantum dots. Attachments have been achieved using a variety of protein designs and binding strategies, including nonspecific adsorption, antibody attachment, and linkage through engineered protein tags (5-13). Recombinant protein design and attachment strategies are crucial experimental parameters, as the display of the motor on the surface significantly impacts the quantitative assessment of motile activity and detection of nm-scale conformational changes that lead to motility (13). Additionally, motor attachment strategies must be orthogonal to immobilization methods of other assay components (e.g., actin) to avoid artifactual binding events. Thus, development of multiple, robust, site-specific attachment strategies for motor proteins is important for the field.

We previously identified a monoclonal antibody (10F12.3) that recognizes a specific epitope in the flexible, coiled-coil, subfragment-2 (S2) domain of a chicken skeletal muscle myosin (13). This monoclonal antibody is ideally suited for coverslip-immobilization of myosin in actin-gliding motility assays (13, 14), as it produces uniform surfaces that support continuous actin gliding while facilitating control of myosin surface density. Here we map the 10F12.3 epitope and determine the sequence (S2Tag) recognized by the mAb. We show that S2Tag can be appended to the S2 domain of β-cardiac heavy meromyosin (β-cHMM), and that purified proteins effectively support surface attachment of β-cHMM to support actin gliding and reproducible optical trap experiments. Additionally, the S2Tag design was used to make a β-cHMM with a human dilated cardiomyopathy (DCM) motor mutation (S532P) that has 100% penetrance and is associated with sudden cardiac death (15). We characterized the effect of this mutation on the actin gliding velocity of the S532P myosin, and we have assessed its mechanical interaction with WT β-cHMM in gliding filament motility with mixed motor surfaces.

## Results and Discussion

### Epitope Mapping for 10F12.3 on Skeletal Muscle Myosin

We identified a novel monoclonal antibody designated 10F12.3 that reacts with a unique site in the S2 domain of the α-helical coiled-coil rod of fast skeletal muscle myosin (13, 16). The antibody is selective for avian fast skeletal muscle myosin, which was useful for isolation of recombinant avian myosin expressed in non-avian cells (17, 18). 10F12.3 has been extensively characterized for its ability to capture and tether native and recombinant avian myosin to surfaces for motor assays (13, 19). We mapped the epitope within the sequence of the myosin S2 domain by combining immunoelectron microscopy, sequence comparison of a family of cross-reacting and non-reacting myosin isozymes, antibody pull-down assays of engineered nested truncations and point mutations of an expressed fragment of the myosin S2 domain (Fig. 1).

**Figure 1.**
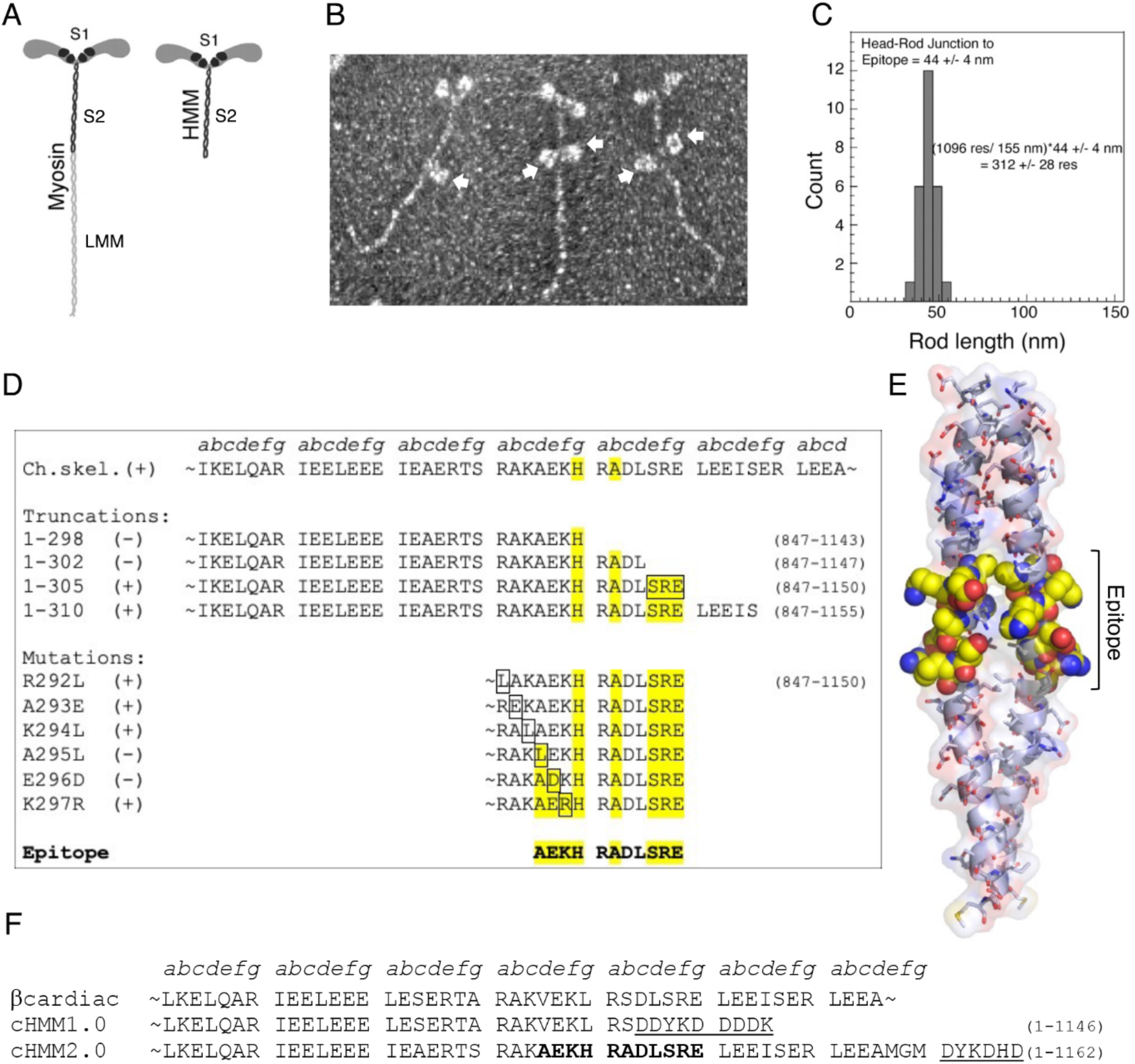
Mapping the epitope recognized by mAb 10F12.3 on skeletal myosin. **A**. Cartoon of the structure of the striated muscle myosin and the HMM subfragment. **B**. rotary shadow electron micrographs of antibody-myosin complexes show the binding of the IgG_1_ mAb 10F12.3 to the S2 domain of the myosin rod. The arrows mark the mAb and >40% of the molecules had two antibodies bound to the rod. **C**. Measurements map the epitope 44 nm from the head-rod junction. The rod is a continuous coiled-coil α-helix of 1096 residues/chain and measures 155 nm long. This yielded an estimate for the location of the epitope of 312 ± 28 residues from the start of the S2 domain. **D**. A series of truncations map the C-terminal limit of the epitope to 305 residues from the start of the S2 domain (the highlighted SRE sequence). The N-terminal limit was determined by the loss of reactivity with the A295L point mutation. This defined the sequence A_1140_EKHRADLSRE_1150_ as the epitope. Residues highlighted in yellow are likely to be key to antibody recognition based on sequences of reactive and non-reactive myosin isoforms (Fig S1). **E**. A model based on the structure of a Lethocerous myosin rod (PDB: 7KOG) flanking this epitope illustrates how the key residues are displayed on the coiled-coil structure. **F**. Sequences showing how the epitope was inserted into the S2 domain of a human β-cardiac myosin for use in motor assays.

Immunoelectron microscopy mapped the 10F12.3 binding site 44 nm from the head-rod junction (Fig 1B, C). This corresponds to 312 ± 28 residues from the start of the S2 domain. A series of S2 fragments truncated at positions ranging from 278-334 amino acids away from the start of the myosin rod were designed for protein expression via an in vitro transcription/translation assay (20). Radiolabeled protein was assayed by immunoprecipitation with mAb 10F12.3 mapping the C-terminal limit of the epitope to 305 residues from the start of the S2 domain (Fig. 1D and Fig. S2). The N-terminal limit was determined by point mutations in residue 292 - 297 of the S2 domain (Fig. 1D). Combining the results of immunoprecipitation of the truncated and mutated S2 segments, together with the pattern of cross reactions with muscle myosin from a variety of species (Fig. S1), the proposed epitope is contained within an 11-residue sequence ‘AEKHRADLSRE’, which we call S2Tag.

### Design and construction of S2Tag containing WT-cHMM

Human β-cardiac myosin has a sequence homologous to chicken skeletal muscle myosin in the S2 domain encompassing the 10F12.3 epitope, but it has substitutions at key residues within the sequence that preclude 10F12.3 binding (Fig. 1F). We have been working with an expressed β-cHMM that included a 42 heptad repeat S2 domain and a C-terminal Flag tag. This construct (β-cHMM1.0) designed before we located the epitope fortuitously interrupted the S2Tag sequence. To demonstrate that the S2Tag sequence encompasses the 10F12.3 epitope we designed a revised version of the human β-cHMM (β-cHMM2.0) to insert the S2Tag, adding two additional heptads of the cardiac S2 sequence after the S2Tag to ensure a stable coiled-coil motif before a C-terminal FLAG sequence (Fig. 1F). The sequence was engineered into an AdEasy shuttle vector for adenovirus production and expression of the β-cHMM2.0 in C2C12 myotubes (11, 21, 22). This shuttle vector was also used to generate a variant containing the DCM mutant S532P as discussed below.

The WT and S532P DCM variants of the β-cHMM2.0 expressed well in C2C12 cells and were readily purified (Fig. S3). Successful antibody-S2Tag binding activity was demonstrated using an antibody capture motility assay (Fig. 2). The 10F12.3 antibody was adsorbed to nitrocellulose-coated glass coverslips, blocked against further protein binding with bovine serum albumin (BSA), and then incubated with purified motor proteins (see Methods). β-cHMM2.0 bound to 10F12.3 mAb coated surfaces and supported smooth gliding of actin filaments (Fig 2 and Fig. S4, (Movie S1)). The proteins do not bind to blocked surfaces lacking the mAb 10F12.3, and the β-cHMM lacking the S2Tag (WT-cHMM1.0) does not bind to the antibody coated surfaces nor support actin movement.

**Figure 2.**
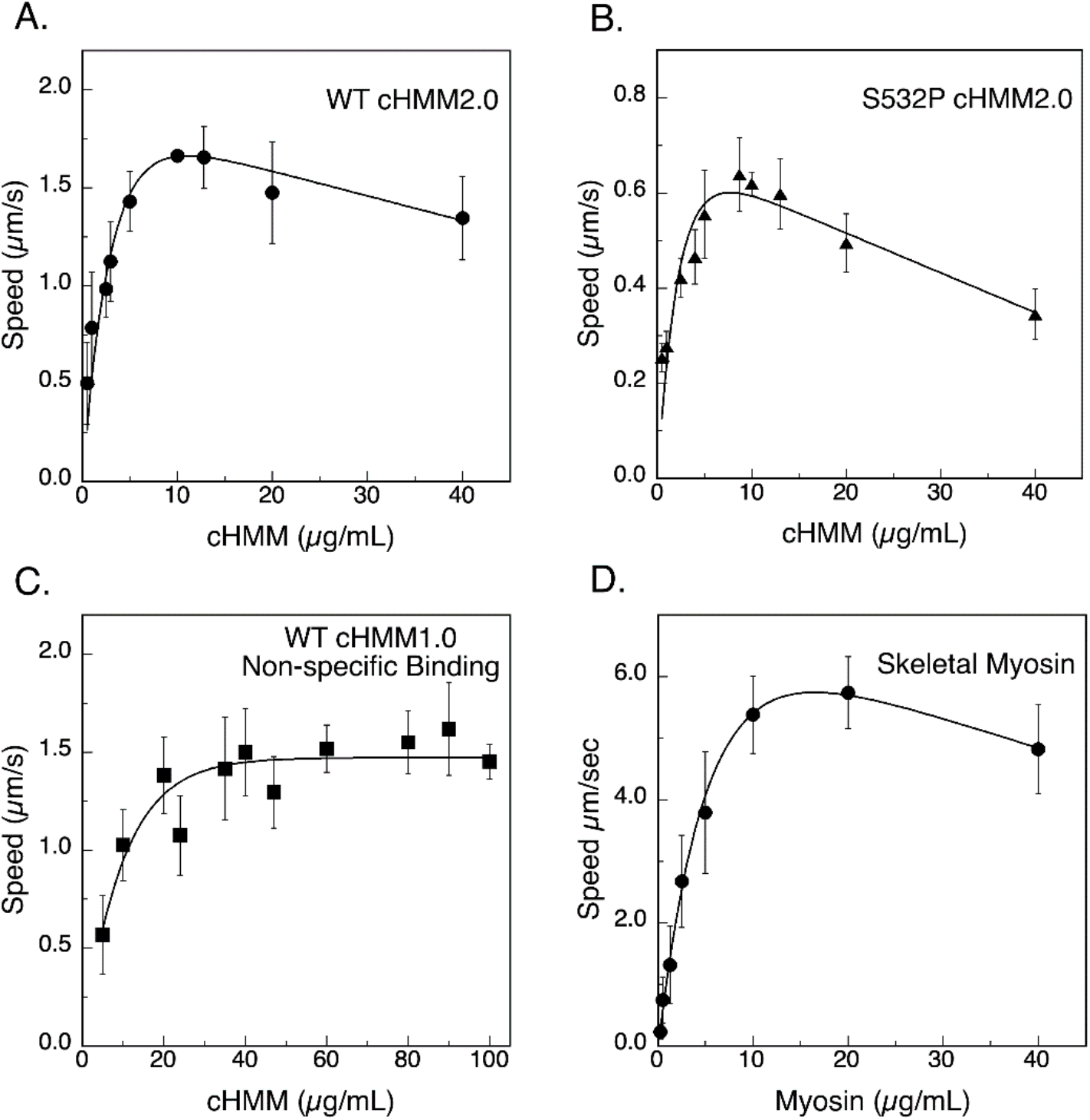
Myosin concentration dependence for motility. An important factor for determining the velocity of actin filament is the myosin surface density. Nitrocellulose coated glass coverslips coated with mAb 10F12.3 were prepared under standard conditions and incubated with increasing concentrations of β-cHMM2.0 and motility recorded and actin gliding speed quantified for: **A**. WT cHMM2.0, and **B**. S532P cHMM2.0. Actin filament speed increases with motor density and is maximal at 5 - 10 µg/mL loading concentration with maximal gliding speed characteristic of the protein (Table 1). **C**. WT cHMM1.0 was bound by non-specific attachment directly to nitrocellulose coated coverslips. Actin filament speed is comparable to antibody captured β-cHMM2.0, but the non-specific binding requires an 8 - 10-fold higher concentrations of myosin to saturate actin filament velocity. **D**. Antibody capture with avian fast skeletal myosin produces maximum actin speed in the same range of loading protein concentration as the β-cHMM2.0 proteins. Motility assays of the β-cHMM proteins were done at 32°C, and the much faster skeletal myosin was measured at 27°C to aid analysis. A smooth curve was used to illustrate the pattern of the data.

Dependence of actin gliding speed on motor surface density (Fig. 2) shows that an optimum speed was achieved in the range of 5 – 10 µg/mL for both the WT and S532P β-cHMM2.0 (Fig. 2 A, B). This concentration range is 8 – 10-fold lower than needed for non-specific, direct binding to nitrocellulose of the WT-cHMM1.0 construct lacking the S2Tag (Fig. 2C). The concentration dependence of motility for the WT-cHMM2.0 is very similar to that of striated skeletal muscle myosin (Sk-myosin), the antigen precursor of mAb10F12.3 (Fig 2D). The similarity in concentration dependence of gliding speed indicates that the S2Tag sequence effectively reconstitutes the 10F12.3 epitope. The titration curves indicate that the S2Tag and 10F12.3 antibody provide an effective way of controlling myosin surface density. We previously found the maximum actin filament speed is achieved at a surface density of ∼600 molecules of myosin/µm^2^, with higher motor densities imparting a drag that slows gliding speed (13, 14, 19).

**Table 1.** Summary of Actin Filament Speeds and Mixed Motor Force Measurements.

| **Unloaded Gliding Actin Filament Velocity** |  |  |  |  |
| --- | --- | --- | --- | --- |
| **Sample** | **Attachment Mode** | **Optimum Concentration $\mu\text{g/mL}$** | **Speed ± SE** | **N** |
| WT β-cHMM2.0 | S2Tag | 10 | $1.69 \pm 0.03$ | 9 |
| WT β-cHMM1.0 | NS | 60 - 80 | $1.49 \pm 0.03$ | 10 |
| S532P β-cHMM2.0 | S2Tag | 10 | $0.62 \pm 0.02$ | 5 |
| Skeletal Myosin | S2Tag | 10 | $6.57 \pm 0.17$ | 4 |
| **Mixed Motor Motility Assay Parameters and Derived Force Ratios** |  |  |  |  |
| **Mixture Slow / Fast** | **$a_s / F_{0s}$** | **$a_f / F_{0f}$** | **$V_{\text{max}s} / V_{\text{max}f}$** | **$F_{0s} / F_{0f}$** |
| S532P / WT | 0.17 | 0.17 | 0.37 | $1.0 \pm 0.1$ |
| WT / Skeletal myosin | 0.17 | 0.25 | 0.25 | $3.4 \pm 0.1$ |
| S532P / Skeletal myosin | 0.17 | 0.25 | 0.09 | $2.9 \pm 0.1$ |

The upper panel lists the actin gliding speed of β-cHMM and Sk myosin measured with the motors attached to 10F12.3 coated surfaces via the S2Tag epitope and binding (S2Tag), or via non-specific (NS) binding directly to nitrocellulose. The optimum motor protein concentration for surface preparation is tabulated along with the average speed ± standard error of the mean for N independent assays.

The gliding speed of the WT β-cHMM2.0 (1.69 µm/s; N = 12) is slightly faster than that of the β-cHMM1.0 non-specifically bound to nitrocellulose (1.49 ± 0.09; N = 10), possibly due to drag imposed by the non-uniform attachment of motors when bound directly to the nitrocellulose (Table 1). The S532P DCM mutation gliding speed (0.62 ± 0.04 µm/s; N = 5) was significantly slower than the WT β-cHMM2.0, consistent with previous reports for this mutation (23). Sk-myosin is a fast muscle myosin with a much higher gliding speed (6.6 ± 0.2 µm/s; N = 4) than β-cardiac myosin, as expected.

### S2Tag Immobilized WT-cHMM in Optical Trapping Assay

Optical trapping enables the measurement of the mechanochemistry of myosin molecules, revealing working stroke displacements on the nanometer scale and actin attachment durations on the millisecond timescale (24, 25). To achieve these resolutions, motors must be bound to surfaces in a way that does not impede function. Although non-specific adsorption to surfaces can be useful, site-specific attachment is preferable, as it results in well-defined and reproducible actomyosin attachments.

To test the utility of the S2Tag in single-molecule studies, we performed a three-bead optical trap assay, in which the pedestal beads were sparsely coated in 10F12.3 antibody and WT-cHMM2.0 was adhered to the antibody rather than directly to the beads (Fig. 3A). An actin filament was suspended between two optically trapped beads to create a ‘dumbbell’ and allowed to interact with single myosin molecules on the pedestal bead in the presence of 1 µM ATP. Imaging chambers in which no 10F12.3 antibody was deposited demonstrated no interactions between the pedestal bead and actin filament. When myosin was adhered *via* the antibody, the dumbbells interacted with the pedestal beads and clear displacements were detected (Fig. 3B). Actomyosin binding interactions were identified by a decrease in the dumbbell bead covariance (22). The attachment durations of these interactions were well fit by a single exponential, with a *k*_detach_ of 5.9 s^-1^, indicating that ATP binding at 1 µM was rate limiting for detachment (Fig. 3C).

**Figure 3.**
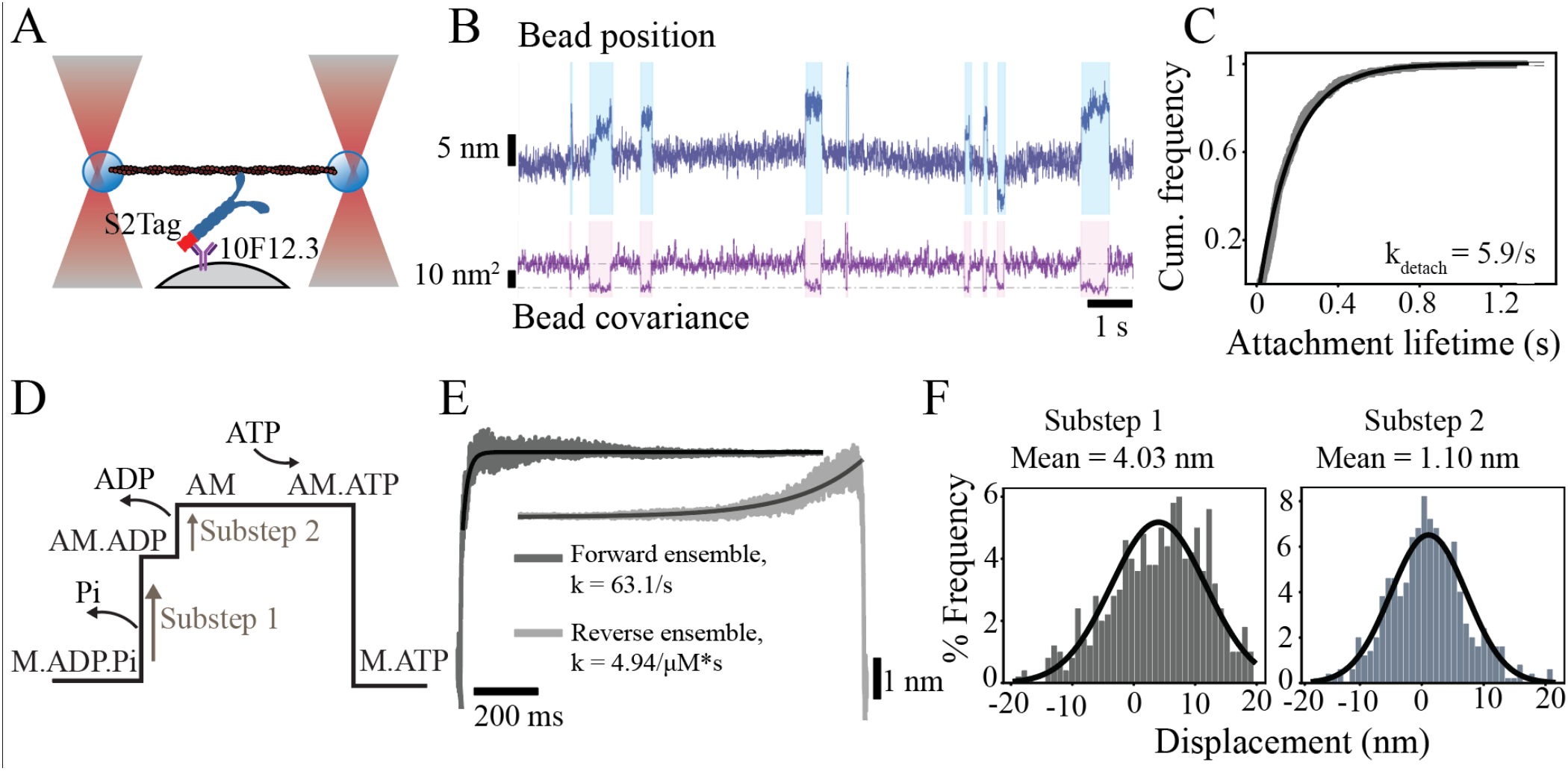
Single-molecule actomyosin interactions of WT-cHMM bound via the S2Tag. **A**. Cartoon of the 3-bead geometry (not to scale). **B**. Sample trace of bead position and covariance for an optically trapped actin dumbbell interacting with WT-cHMM2.0 adhered to the pedestal bead through the 10F12.3 antibody-epitope interaction. **C**. Cumulative distribution of attachment durations of actomyosin interactions. **D**. Schematic of the two-substep power stroke of cHMM and associated product release. **E**. Ensemble averages of actomyosin interactions aligned at start (Forward ensemble) or end (Reverse ensemble). **F**. Histograms of substep sizes for single interactions.

The β-CM power stroke occurs in two substeps, the first associated with phosphate release and the second with ADP release, while actin detachment follows ATP binding (Fig. 3D; (26)). By aligning the events at the onset of interactions and averaging forward in time, or by aligning at the end and averaging backward in time, we generated ensemble averages of single-molecule interactions, which demonstrated a clear two-substep power stroke (Fig. 3E). The transition rate from state 1 to state 2 occurred at a rate of 63.1 s^-1^, consistent with the rate constant for ADP release of WT-cHMM (21). The transition from state 2 to detachment occurred at a rate of 4.94 s^-1^, consistent with the second-order ATP binding rate of 4.6 μM^-1^s^-1^ (21). The displacements associated with each event’s substeps approximate a normal distribution with a mean substep 1 displacement of 4.03 nm and a mean substep 2 displacement of 1.10 nm, both consistent with previously published values for WT-cHMM (Fig. 3E; (22, 26)). Thus, the 10F12.3 epitope-antibody interaction provides a robust method for adhesion of cHMM to pedestal beads for single-molecule studies; indeed, in our hands, this adhesion scheme significantly reduced the variability of loading concentrations required to achieve single-molecule interactions compared with non-specific adhesion.

### Mechanical interactions between WT-cHMM and S532P-cHMM and mutant cardiac myosin

Precise control of myosin attachments to the motility surface makes it possible to assess mechanical interactions between different myosin paralogs or between wild-type and mutant motors using mixed motility assays (27, 28). When two myosins with different cycling rates bind to and participate in moving the same actin filament, the two motors mechanically interact through the filament to determine filament velocity. A key to detecting the interaction is a significant difference in gliding filament speed so one can compare a faster myosin to a slower myosin. The slower myosin imposes an internal load against which the faster cycling myosin must act. If the motors have similar duty ratios, similar generated force, and force-dependence of actin detachment rates (i.e., similarly shaped force-velocity (F-V) curves), then the filament velocity will be linearly related to the fraction of each species. However, if the forces are not balanced and the other parameters are similar, the stronger motor overpowers the weaker motor, resulting in non-linear dependence on the fraction of each species ((27, 28); see Methods).

To assess the utility of the S2Tag in mixed motility assays, we performed experiments with the WT and S532P β-cHMM2.0 mixed with each other and each mixed with Sk-myosin. The biochemical kinetics of myosin with the S532P mutation were characterized previously (23) and showed a reduced steady-state actin-activated ATPase rate (3-fold) and ∼3-fold slower gliding speed consistent with our observations (Fig. 2) (23).

Actin gliding speeds were measured with pairwise mixtures of WT-cHMM2.0, S532P-cHMM2, and Sk-myosin all bound to 10F12.2 coated coverslips (Fig. 4). The speed of the mixed WT-cHMM2.0 and S532P-cHMM2.0 motility increased linearly with increased fraction of WT-cHMM2.0 (Fig. 4A), suggesting that the force generated by S532P-cHMM is comparable to the WT protein, despite its slower kinetics. In contrast, both WT- and S532P-cHMM2.0 showed concave-upward dependencies of the actin gliding speeds when mixed with the much faster Sk-myosin (Fig. 4C, D). The slower cardiac myosins strongly impact the gliding speed of the faster skeletal myosin. Assuming similarly shaped F-V curves, this suggests that they generate more force than the fast skeletal myosin. Using the model equations from Harris and Warshaw (27, 29), the fitted values for the apparent force ratio of the slower to faster myosins (*F*_*os*_ */F*_*of*_) suggest the cardiac proteins are both ∼3 fold stronger than the Sk-myosin (Table 1). The Sk-myosin in this instance also acts as a reference for confirming that the WT and S532P β-CM produce comparable forces but with distinct kinetics.

**Figure 4.**
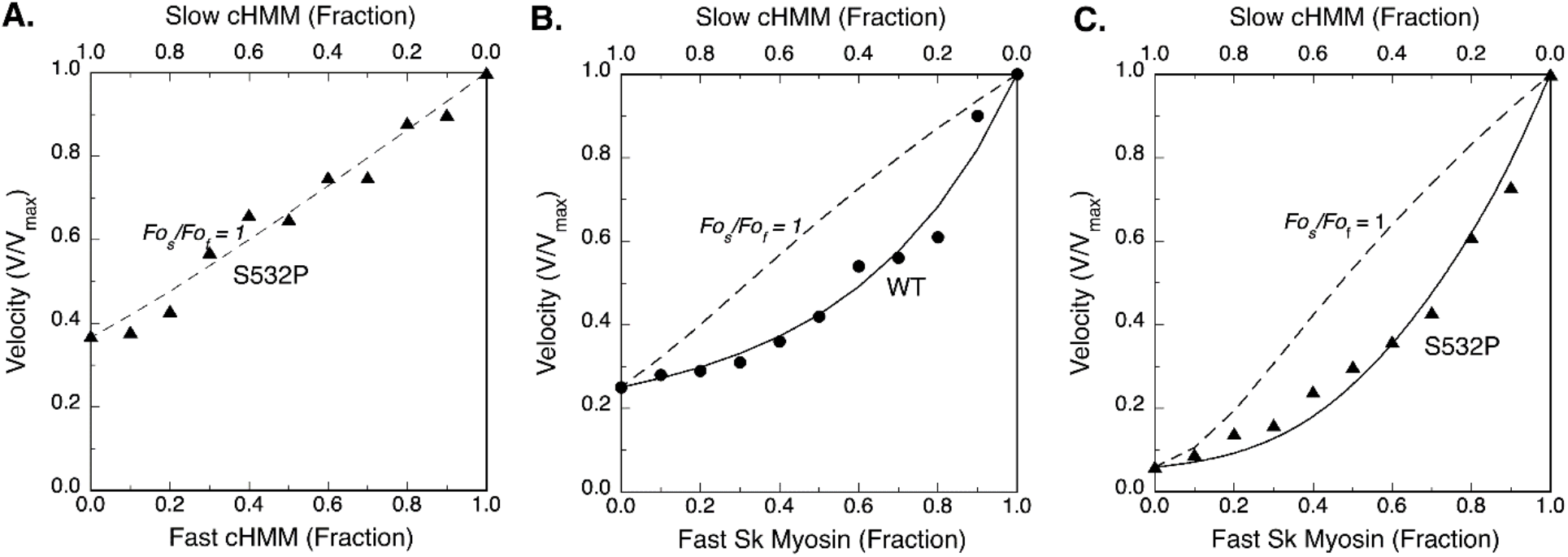
Mixed myosin motility assay. **A**. Assay of WT cHMM2.0 (fast myosin) mixed with S532P cHMM2.0 (slow myosin) producing a linear relationship between actin filament velocity and the fraction of fast myosin consistent with a unitary force ratio, 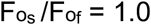. The dotted line in each figure is the calculated fit if the force produced by each isoform is equal. Despite having a slower speed, S532P cHMM2.0 apparently produces force comparable to WT cHMM2.0. **B**. Avian skeletal muscle myosin is a very much faster myosin than the β-CM, and the concave fit when assayed with the slower WT cHMM2.0 yields a force ratio, 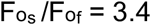, suggesting the slower cardiac myosin generates higher force. **C**. The S532P mutation imparts about the same load on the skeletal myosin, 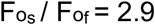, as the WT cHMM2.0. This result is consistent with balanced forces observed in the direct comparison of WT and S532P cHMM2.0 (Panel A).

## Conclusion

We have described and mapped a unique epitope in a conserved region of striated myosin S2. The antibody-antigen method of attachment of myosin to surfaces allows for reproducible control of the motor surface density that optimizes the unloaded gliding of actin filaments *in vitro*. The uniform attachment of the myosin has been shown to optimize actin filament speed, to highlight the mechanical interaction between different isoforms, and to minimize contributions from sub-optimal interactions from non-specific modes of attachment at an actin reaching distance of about 62 nm from the surface (13). Analysis of a cardiac myosin mutation, S532P, that causes DCM, shows the power of assessing the impact of the cardiomyopathy mutation on the activity of the WT β-cardiac myosin. The method of attaching molecular motors to a microscope slide for functional assays improves reliability and reproducibility of experiments and should facilitate further studies of myosin and other molecular motors.

## Experimental Procedures

### Proteins

Myosin was prepared from adult White Leghorn chicken pectoralis muscle as described previously (30). Actin was extracted from rabbit skeletal muscle acetone powder (Pel-Freez, Rogers, AK). Immunoelectron microscopy of rotary-shadowed molecules was performed as described previously (30). Monoclonal antibody, 10F12.3, was prepared and characterized as described previously (16, 30). Myosin isoform specificity was determined by enzyme linked immunoassays and Western blotting; the mAb 10F12.3 reacts with embryonic, post-hatch, and adult fast skeletal muscle myosin from chickens. It does not cross-react with rabbit myosin. IgG_1_ class monoclonal antibodies were purified on protein A-Sepharose from ascites fluid obtained by passage of hybridoma lines through CAF1/J mice.

### Coupled Translation/Transcription Assay and Immunoprecipitation

Coupled transcription and translations were performed with TNT Quick kits purchased from Promega (Madison, WI), and supplemented with Redivue L-[^35^S] Methionine (GE Healthcare). The myosin S2 expression vectors used in coupled translation assays contain the coding sequences downstream of an SP6 promoter in a pGEM4 vector. Coupled transcription-translation assays were incubated 2 hr at 30°C with 0.5 mg plasmid DNA per 50 mL reaction (31). Construction of the vectors for the embryonic chicken skeletal muscle S2 has been described in detail elsewhere (32). An aliquot of the lysate was incubated with 5 µg of the anti-S2 IgG antibody for 2 hr at 4°C. Protein A agarose beads (1:1 suspension) were added and incubated for 1 hr at 4°C. The Protein A beads were washed with 1 mL of 10 mM Imidazole, 150 mM NaCl, 0.05% NP-40 for 30 min and twice for 10 min at 4°C on a rotating rocker followed by a final wash in the same buffer without detergent. Bound proteins were eluted into SDS-PAGE gel loading buffer and analyzed by SDS-PAGE and autoradiography.

### Adenovirus manipulation

Our original human β-cardiac HMM (cHMM) design encodes residues 1-1137 of the *MYH7* gene (GenBank: AAA51837.1) with a FLAG tag added at the C-terminus (1138-1146) of the S2 domain (cHMM1.0) (21, 22).The cHMM cDNA was cloned into the pShuttle-IRES-hrGFP-1 vector (Agilent Tech., Santa Clara, CA). The AdcHMM-Flag virus was prepared and amplified for expression of cHMM protein in C2C12 cells (17, 33). For the cHMM2.0 construct, the sequence of the epitope (AEKHRADLSRE) was introduced into the coiled-coil S2 domain of β-cHMM, followed by two additional heptads of cardiac S2 sequence and a FLAG tag at the C-terminus based on the sequence in Fig. 1. The new DNA sequence was constructed by GeneWiz (Azenta Life Sciences, South Plainfield, NJ) and inserted into the original AdEasy shuttle vector for adenovirus production and sequenced. The WT shuttle vector (pSH-cHMM2.0-FLAG-hrGFP) was then used to generate the β-cHMM mutant protein containing the DCM mutation, S532P (GeneWiz). The adenovirus plasmids were prepared and sequenced before new virus stocks were isolated and amplified in Ad293 cells through 5 passages to produce high titer virus stocks. The virus was harvested and purified by CsCl density sedimentation yielding final virus titers of ∼10^11^ plaque forming units per mL (pfu·mL^−1^) for infection of C2C12 cells and protein production.

### Muscle cell expression and purification of β-cardiac HMM

Maintenance of the mouse myogenic cell line, C2C12 (CRL 1772; American Type Culture Collection, Rockville, MD), has been described in detail elsewhere (17). Confluent C2C12 myoblasts were infected with replication defective recombinant adenovirus (AdcHMM2.0) at 2.7 X 10^8^ pfu·mL^−1^ in fusion medium (89% DMEM, 10% horse serum, 1% FBS). Expression of recombinant cHMM was monitored by accumulation of co-expressed GFP fluorescence in infected cells. Myocyte differentiation and GFP accumulation were monitored for 216 – 264 hr after which the cells were harvested. Cells were chilled, media removed, and the cell layer was rinsed with cold PBS. The cell layer was scraped into Triton extraction buffer: 100 mM NaCl, 0.5% Triton X-100, 10 mM Imidazole pH 7.0, 1 mM DTT, 5 mM MgATP, and protease inhibitor cocktail (Sigma-Aldrich, St. Louis). The cell suspension was collected in an ice-cold Dounce homogenizer and lysed with 15 strokes of the tight pestle. The cell debris in the whole cell lysate was pelleted by centrifugation at 17,000 x g for 15 min at 4°C. The Triton soluble extract was fractionated by ammonium sulfate precipitation using sequential steps of 0-30% saturation and 30-60% saturation. The cHMM precipitates between 30-60% saturation of ammonium sulfate. The recovered pellet was dissolved in and dialyzed against 50 mM Tris, 150 mM NaCl, pH 7.4, 0.5 mM MgATP for affinity purification of the FLAG-tagged cHMM on M2 mAb-Sepharose beads (Sigma-Aldrich). Bound cHMM was eluted with 0.1 mg·mL^−1^ FLAG peptide (Sigma-Aldrich). Protein was concentrated and buffer exchanged on Amicon Ultracel-10K centrifugal filters (Millipore; Darmstadt, Germany), dialyzed exhaustively into 10 mM MOPS, 100 mM KCl, 1 mM DTT before a final centrifugation at 300,000 x g for 10 min at 4°C. Aliquots were drop frozen in liquid nitrogen and stored in vapor phase at –147°C.

### *In vitro* gliding filament motility assay

Measurement of *in vitro* motility of human β-cHMM2.0 was done as previously described for skeletal muscle myosin (11, 13, 21, 34). Nitrocellulose-coated glass coverslips were incubated with 0.15 mg/mL of the mAb, 10F12.3, followed by blocking the surface with 1% BSA. β-cHMM2.0 proteins were diluted in motility buffer (MB) (25 mM imidazole, pH 7.8, 25 mM KCl, 4 mM MgCl_2_, 1 mM MgATP, 1 mM DTT) supplemented with 1% BSA (MB/BSA) to the final concentration as required. The antibody-coated coverslips were incubated with β-cHMM2.0 for ∼2 hr in a humidified chamber at 4 °C. The coverslips were washed with MB/BSA, followed by actin blocking with 1 μM F-actin, and washes with motility buffer, then transferred to a 15-μL drop of 2 nM rhodamine-phalloidin–labeled actin in a modified motility buffer (with 7.6 mM MgATP, 50 mM DTT, 0.5% methyl cellulose, 0.1 mg/mL glucose oxidase, 0.018 mg/mL catalase, 2.3 mg/mL glucose) in a small parafilm ring fixed on an alumina slide with vacuum grease. The chamber was observed with a temperature-controlled stage and objective set at 32 °C on an upright microscope with an image-intensified charge-coupled camera capturing data to an acquisition computer at 5–30 fps. depending on assay parameters. Movement of actin filaments from 500 to 1,000 frames of continuous imaging was analyzed with semi-automated filament tracking programs as previously described (34). The trajectory of every filament with a lifetime of at least 10 frames was determined; the instantaneous velocity of the filament moving along the trajectory, the filament length, the distance of continuous motion and the duration of pauses were tabulated. A weighted probability of the actin filament velocity for hundreds of events was fit to a Gaussian distribution and reported as a mean velocity and SD for each experimental condition.

### Three-bead optical trap assay

Flow cells were constructed on a microscope slide (Corning), with double-sided tape adhering a coverslip coated with 0.1% nitrocellulose (Electron Microscopy Services) mixed with 2.5-µm-diameter silica beads. 10F12.3 mAb at 0.03 mg/mL in trapping buffer (25 mM KCl, 60 mM MOPS pH 7.0, 1 mM DTT, 1 mM MgCl_2_, 1 mM EGTA) was incubated for 6 s, followed by blocking twice with 1 mg/mL BSA in trapping buffer for 3 min each. S2tag-myosin, diluted to 1 µg/mL in myosin buffer (trapping buffer with 300 mM KCl), was added to the chamber and incubated for 3 min. Two blocking steps with 1 mg/mL BSA were repeated for 2 min each. Finally, trapping buffer was added with 1 µM MgATP (determined spectroscopically at 259 nm with extinction coefficient of 15.4 mM^-1^*cm^-1^), 0.2 nM rabbit skeletal muscle actin filaments with 10% biotinylated actin stabilized by rhodamine-phalloidin at a 1.1 molar ratio with actin monomers, 2.5 mg/mL of glucose, and fresh glucose oxidase + catalase mix (Sigma). Finally, 0.4 ng of 750 nm-diameter polystyrene beads (Polysciences, Warrington, PA), which had been incubated overnight at 4°C with rotation in 10 mg/mL neutravidin, was added to one side of the chamber, and the chamber was sealed with vacuum grease.

Optical trapping was performed as previously described in a laboratory-built dual-beam optical trap (35). An actin dumbbell was positioned to maximize actomyosin displacement and actomyosin interactions were detected by a decrease in bead-bead covariance. Interactions were detected by a drop in bead covariance. All molecules analyzed had >75 individual interaction events with a single actin dumbbell. Ensemble averages were generated as described previously (22). Plots were fit by single exponential functions. Step sizes were measured as described previously (22).

### Derivation of relative force in mixed myosin motility assay

The analysis of actin filament sliding velocity over mixtures of myosin isoforms of differing unloaded shortening velocity and/or force production is based on the approach developed by the Warshaw group (27, 29). Briefly, if two myosin isoforms with differing cycling rates are arranged such that they can interact simultaneously with the same actin filament, then the actin filament sliding velocity will reflect the mechanical interaction between the different myosin isoforms. The assay design requires two myosin isoforms that differ sufficiently in unloaded shortening velocity to be distinguishable. The myosin molecules are randomly arrayed on a surface so that sliding movement of actin is driven by both isoforms. In the assays described here a surface is prepared with a mAb and both myosin isoforms bind to the same mAb via their common epitope. The proportion of slow/fast myosin is varied during preparation of the surfaces and the resultant actin filament gliding velocity measured with the motility assay.

The approach assumes that both positive forces and resisting compression forces vary with isoform ratio and impact actin filament velocity, and that the force-velocity curve for the individual myosin isoforms has the same curvature (*a*/*F*_*o*_) as force-velocity relationship of the muscle from which the myosin has been isolated (28). These approximations have been validated for a variety of myosin isoforms including cardiac and skeletal muscle myosin and this assay has provided a method to rank relative force production for these myosin isoforms (9, 27, 28, 36).

The change of actin filament velocity (V_actin_) as a function of slow/fast isoform ratio has been modeled mathematically as a quadratic function of actin filament velocity (V_actin_) as follows:

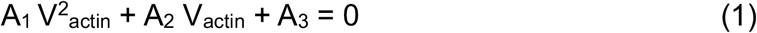

The constants A_1_, A_2_, and A_3_,

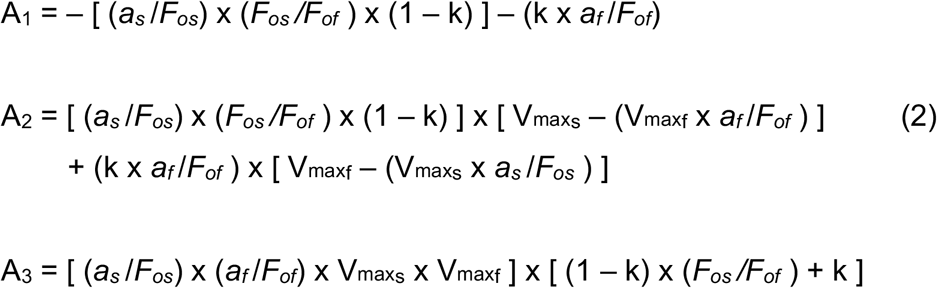

are defined by the fraction of the faster (k) and slower (1-k) myosin, the hyperbolic constants for the force-velocity curves of the corresponding slower (*a*_*s*_ /*F*_*os*_) and faster (*a*_*f*_ /*F*_*of*_) muscles, and the maximum actin filament velocity for the slower 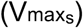 and faster 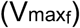 myosin. Values assumed from skeletal and cardiac muscle and fixed in the analysis here were *a*_*f*_ /*F*_*of*_ = 0.25 and *a*_*s*_ /*F*_*os*_ = 0.17, respectively.

## Data Availability

Representative plots for all experimental data and unedited SDS-PAGE/coomassie stained gels and autoradiographs are presented in the manuscript and supplement. All fits and statistics are reported. Raw data is available upon request from D.A.W. The hybridoma cell line 10F12.3 is being prepared for deposit in the Developmental Studies Hybridoma Bank, University of Iowa.

## Author Contributions

Experiments were conceptualized by D.A.W, B.B, E.M.O and R.C.C and Y.E.G. Viruses, antibodies, vectors, and all other proteins were prepared by D.A.W., B.B. and R.C.C. Experimental work was done by B.B., R.C.C. and D.A.W. The manuscript was written by B.B., D.A.W, R.C.C., E.M.O, and Y.E.G.

## Supporting Information

This article contains supplemental information.

## Competing Interests

The authors declare no competing interests.

## Acknowledgements

The authors acknowledge the assistance of Jie Liu in the early stages of the epitope mapping. The work was supported by PHS grants R01 HL157997 to E.M.O, Y.E.G., and D.A.W., R35 GM118139 to Y.E.G., R37 GM057247 to E.M.O., and NSF grant (CMMI:15-48571) to Y.E.G. and E.M.O.

## Notes

### Competing Interest Statement

The authors have declared no competing interest.

