## Supplemental Figures for "S2Tag, a novel affinity tag for the capture and immobilization of coiled-coil proteins: application to the study of human β-cardiac myosin"

[1]Department of Pathology and Laboratory Medicine, Robert Wood Johnson Medical School, Rutgers University, Piscataway, NJ 08854, [2]Department of Physiology and Pennsylvania Muscle Institute, Perelman School of Medicine, University of Pennsylvania, Philadelphia, PA 19104, and [3]Department of Pharmacology and Department of Molecular and Cellular Biology, University of California at Davis, Davis, CA 95616

Corresponding author:
Donald A. Winkelmann

Department of Pathology and Laboratory Medicine, Robert Wood Johnson Medical School, Rutgers University, Piscataway, NJ 08854

**This document contains Supplementary Materials:**

- Figures S1-S4
- Movie S1

A

```
                                                                                  EM
             1109                                                                 ↓ | Weak Point |                            1192
Heptad       abcdefg abcdefg abcdefg abcdefg abcdefg abcdefg abcdefg abcdefg abcdefg abcdefg abcdefg abcdefg

ChEmMyob     gm----- ------- ------- -----t- ------h -a----- ------- ------- -at--dm ------- ---m--- -------
ChEmMyod     gm----- ------- t------ -----t- ------h -a----- ------- ------- -aa--dm ------- ---m--- -------
ChMyN116     gm----- ------- ------- -----t- ------h -a----- ------- ------- -aa--em ------- ---m--- -------
ChMyN118     gm----- ------- ------- -----t- ------h -a----- ------- ------- -aa--dm ------- ---m--- -------
RatEmMyo     sl----- ------- ------- -----at ---t--q -s-ya-- ---l--- ------v -st--el ------- -l-l--- -------
HumEmMyo     gl-f--- ------- ------- -----at ---t--q -s-ya-- ---l--- ------v -st--el ------- -l-l--- -------
RabFSkMy     am----- ------- ------- -----a- ------q -s----- ------- ------- -sa--em ------- ---m--- -------
HumPnMyo     ei----- ------- ----g-- -----a- ------q -s----- ------- ------- -sa-vel ------- ---l--- -------

Consensus    --QLQKK IKELQAR IEELEEE IEAER-S RAKAEKh RaDLSRE LEEISER LEEAGGA T--QI-- NKKREAE FQK-RRD LEEATLQ
                                                AEKh RaDLSRE
                                                10F12.3 epitope
```

B

```
Heptad       abcdefg abcdefg abcdefg abcdefg abcdefg abcdefg abcdefg abcdefg abcdefg abcdefg abcdefg abcdefg

ChSkel       GMQLQKK IKELQAR IEELEEE IEAERTS RAKAEKH RADLSRE LEEISER LEEAGGA TAAQIDM NKKREAE FQKMRRD LEEATLQ  1192
L.indic.     VSKTQKQ IKELQSR IEELEEE VEAERQA RGKAEKQ RADLARE LEELGER LEEAGGA TSAQIEL NKKREAE MSKLRRD LEESNIQ  1192
HumSkel      GMQLQKK IKELQAR IEELEEE IEAERAS RAKAEKQ RSDLSRE LEEISER LEEAGGA TSAQIEM NKKREAE FQKMRRD LEEATLQ  1188
HumβCard     GSQLQKK LKELQAR IEELEEE LEAERTA RAKVEKL RSDLSRE LEEISER LEEAGGA TSVQIEM NKKREAE FQKMRRD LEEATLQ  1184
HumαCard     ALQLQKK LKENQAR IEELEEE LEAERTA RAKVEKL RSDLSRE LEEISER LEEAGGA TSVQIEM NKKREAE FQKMRRD LEEATLQ  1186
HumSm2B      KNNALKK IRELEGH ISDLQED LDSERAA RNKAEKQ KRDLGEE LEALKTE LEDTLDS TATQQEL RAKREQE VTVLKKA LDEETRS  1196
HumNMIIA     KNMALKK IRELESQ ISELQED LESERAS RNKAEKQ KRDLGEE LEALKTE LEDTLDS TAAQQEL RSKREQE VNILKKT LEEEAKT  1182
HumNMIIB     KNNALKV VRELQAQ IAELQED FESEKAS RNKAEKQ KRDLSEE LEALKTE LEDTLDT TAAQQEL RTKREQE VAELKKA LEEETKN  1220
HumNMIIC     RAQLLKS LREAQAA LAEAQED LESERVA RTKAEKQ RRDLGEE LEALRGE LEDTLDS TNAQQEL RSKREQE VTELKKT LEEETRI  1214
```

**Figure S1**. Myosin S2 sequence comparisons. **A**. Sequence comparison of a family of myosin isozymes that have been tested for cross reaction with mAb 10F12.3. Chicken fast skeletal muscle myosin isozymes (boxed) all cross-react with 10F12.3. Embryonic rat (RatEmMyo) and human (HumEmMyo), rabbit psoas (RabFSkMyo) and perinatal human (HumPnMyo) myosins do not cross-react with 10F12.3. The sequences flanking the epitope defined by immunoelectron microscopy are highly homologous. **B**. Sequence comparison of myosin II proteins in the S2 region flanking the mAb 10F12.3 epitope. The sequence $A_{1140}EKHRADLSRE_{1150}$ that was identified as the epitope in chicken skeletal myosin, can be transferred to other myosin II proteins as shown for β-cardiac myosin in the present study. The myosin sequences are chicken skeletal muscle (NP_989559), *Lethocerus indicus* flight muscle (ASP18627), human skeletal muscle (NP_005954), human β-cardiac (NP_000248), human α-cardiac (NP_002462), human smooth muscle (NP_001035202), human nonmuscle IIA (NP_002464), human nonmuscle IIB (NP_001242941), and human nonmuscle IIC (NP_1070654). Residues highlighted in yellow are key to antibody recognition.

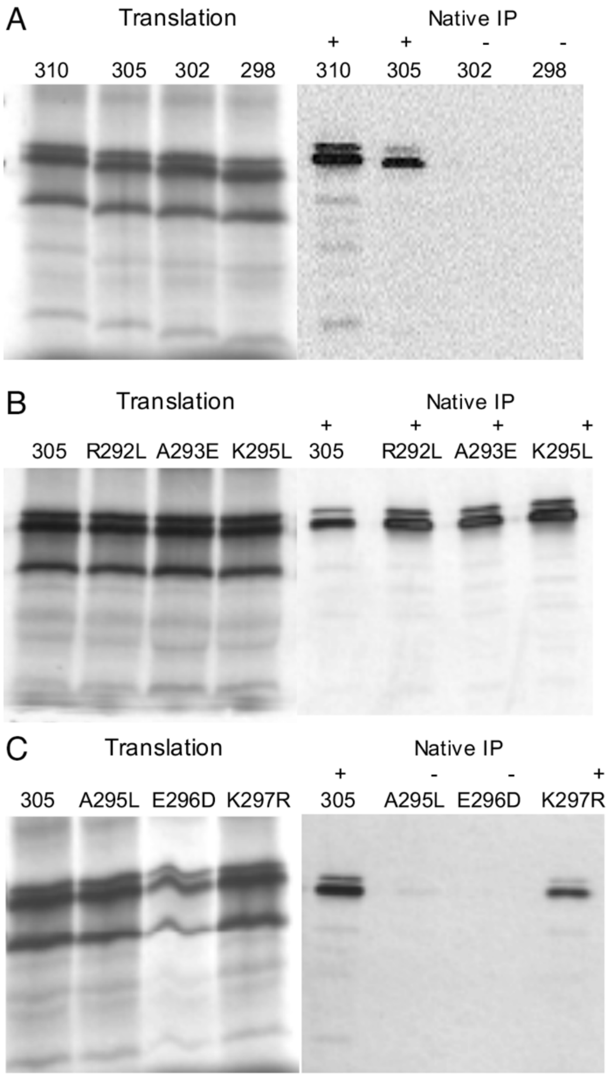


**Figure S2**. Expression and immunoprecipitation of S2 fragments with mAb 10F12.3. A series of S2 fragment truncations and point mutations were expressed in an *in vitro* transcription-translation assay and radiolabeled protein was assayed by immunoprecipitation with 10F12.3. **A.** S2 fragments truncated at positions ranging from 278-334 amino acids downstream from the start of the myosin rod (Fig. 1D) mapped the C-terminal limit of the epitope to residue 305 from the start of the S2 domain. **B, C.** The N-terminal limit of the epitope was determined by site-directed mutagenesis of residues 292 - 297 in the S2 domain truncated at residue 305. Residues A295 and E296 are shown to be key to antibody recognition.

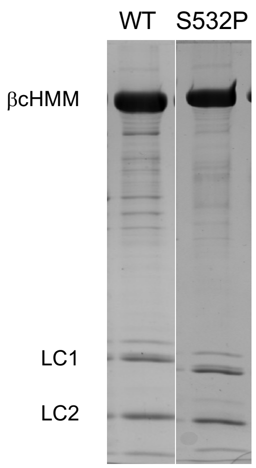


**Figure S3**. SDS polyacrylamide gel electrophoresis of typical purified WT- and S532P-cHMM2.0 protein samples used in this study. Proteins were expressed in C2C12 myotubes via adenovirus mediated delivery of a β-cHMM2.0 expression cassette (see Material and Methods). The purified β-cHMM2.0 has a 134 kDa heavy chain and associated essential and regulatory light chains. There is no detectable C2C12 myosin contamination in the samples.

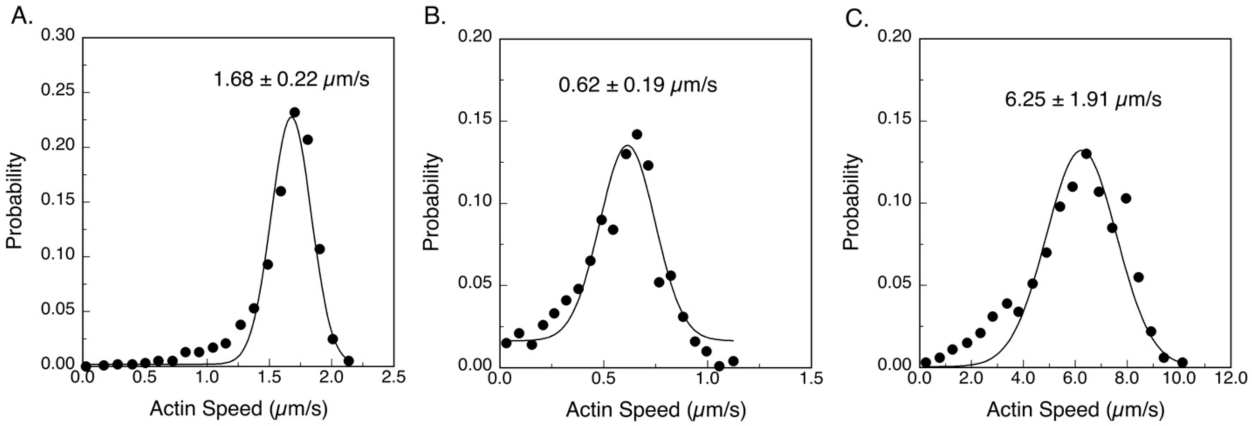


**Figure S4**. Probability distributions of filament speeds from motility assay movies. The data were fit to Gaussian distributions to determine a mean speed and standard deviation for: **A.** WT cHMM2.0, **B.** S532P cHMM2.0, and **C.** Sk-myosin. All proteins were bound to nitrocellulose coverslips coated with the mAb 10F12.3. Actin filament movement was analyzed in movies of 500 - 1,000 frames with a semi-automated filament tracking program. The velocity of the filament centroid moving on the trajectory, the filament length, the distance of continuous motion, and the duration of pauses were tabulated. A weighted probability of the actin filament velocity for hundreds of events were fitted by a Gaussian distribution and reported as a mean velocity and SD for each experimental condition. The data shown here were collected at a motor protein loading concentration of 10 μg/mL and assayed at 32°C.

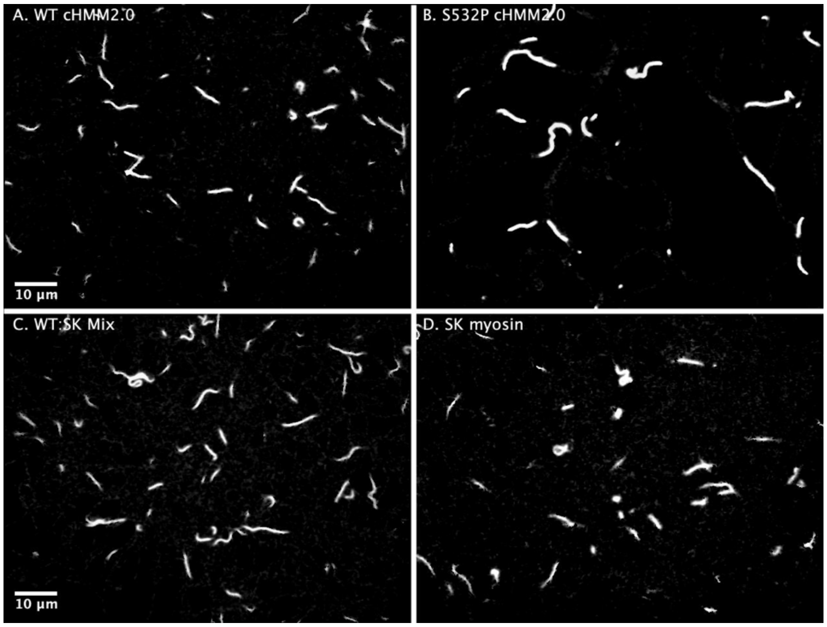


**Movie S1**. Motility assays for **A.** WT-cHMM2.0, **B.** S532P-cHMM2.0, **C.** WT:Sk-myosin mix (50:50), and **D.** Sk-myosin bound to nitrocellulose coverslips coated with mAb 10F12.3. Gliding actin filament movement was recorded at 5 fps for the slower motors: WT-cHMM2.0 (1.68 µm/s), and the S532P mutant (0.62 µm/s). The playback speed for these two panels is 3X faster than the recorded speed. The faster gliding speed of actin moving over the 50:50 mixture of WT-cHMM2.0 with Sk-myosin (3.0 µm/s), and Sk-myosin along (6.3 µm/s) were recorded and played back at 15 fps. All coverslips were prepared at 10 µg/mL motor protein loading. The 500-frame movies shown here were collected at 32°C and are representative of the data presented in Figures 2, 4, and S4 and tabulated in Table 1.
